# RRBP1 rewires cisplatin resistance in Oral Squamous Cell Carcinoma by regulating YAP-1

**DOI:** 10.1101/2020.03.18.998070

**Authors:** Omprakash Shriwas, Rakesh Arya, Sibasish Mohanty, Sugandh Kumar, Rachna Rath, Sandeep Rai Kaushik, Mukund Chandra Thakur, Falak Pahwa, Saroj Kumar Das Majumdar, Dillip Kumar Muduly, Ranjan K Nanda, Rupesh Dash

**Author notes:** Corresponding authors Rupesh Dash, Institute of Life Sciences, Nalco Square, Chandrasekharpur, Bhubaneswar-751023, Odisha, India., and/or Ranjan Nanda, Group Leader, Translational Health Group, International Centre for Genetic Engineering and Biotechnology, New Delhi-110067, India.

## Abstract

Cisplatin-based chemotherapy still remains as one of the primary treatment modalities for OSCC. Several OSCC patients experience relapse owing to development of chemoresistance. To identify key resistance triggering molecules, we performed global proteomic profiling of human OSCC lines presenting with sensitive, early and late cisplatin resistance patterns. From the proteomic profiling study, human RRBP1 was identified to be upregulated in both early and late cisplatin-resistant cells with respect to the sensitive counterpart. Analysis of OSCC patient sample indicates that RRBP1 expression is elevated in chemotherapy-non-responder tumors as compared to chemotherapy-naïve tumors. Knocking out RRBP1 resulted in restoring cisplatin mediated cell death in chemoresistant lines and patient derived cells (PDC). Mechanistically, RRBP1 regulates YAP-1 to induce chemoresistance in OSCC. The chemoresistant PDC xenograft data suggests that knock out of RRBP1 induces cisplatin mediated cell death and facilitates a significant reduction of tumor burden. We also found Radezolid, a novel oxazolidinone antibiotic represses the expression of RRBP1 and restores cisplatin-induced cell death in chemoresistant OSCC. This unique combinatorial approach needs further clinical investigation to target advanced OSCC. Here with for the first time, we uncover the novel role of RRBP1 as potential modulator of cisplatin resistance in advanced OSCC.

## Introduction

HNSCC is the sixth most common cancer globally and most prevalent in developing countries. OSCC is an aggressive form of HNSCC and is the most common cancer among Indian males [1]. Approximately 80,000 new OSCC cases are reported annually with a mortality of 46,000 individuals each year in India. The traditional treatment modalities for advanced OSCC comprises of surgical removal of primary tumors followed by concomitant adjuvant chemoradiotherapy [2]. In addition, neoadjuvant chemotherapy is frequently recommended for surgically unresectable OSCC tumors that reduces tumor and provides more surgical options. Despite having these solutions, the 5-year survival rate of advance tongue OSCC is approximately 50%, indicating a possible resistance to currently available therapeutics.

Chemoresistance is one of the important factors for treatment failure in OSCC [3]. Cisplatin alone or in combination with 5-fluorouracil and taxane/docetaxel (TPF) are generally used as chemotherapy regimen for OSCC [4]. But due to chemoresistance development, patients experience relapse which leads to continued tumor growth and metastasis. The chemoresistant properties could be attributed to enhanced cancer stem cell population, decreased drug accumulation, reduced drug-target interaction, reduced apoptotic response and enhanced autophagic activities [5]. These hallmarks present the endpoint events, when cancer cell had already acquired chemoresistance. Till date, the causative factors responsible for acquiring chemoresistance in cells are yet not explored. Identifying these molecular triggers will enable us to understand the molecular mechanism behind chemoresistance and may be useful to identify important targets.

In the present study, to identify the causative factors responsible for cisplatin resistance, we employed a global quantitative proteomics study to identify deregulated proteins in cisplatin-resistant OSCC cancer cell lines. Protein samples extracted from sensitive, early and late cisplatin resistant cells were subjected to isobaric tags for relative and absolute quantification (iTRAQ) studies using liquid chromatography-tandem mass spectrometry (LC-MS/MS). A representative deregulated protein was selected for validation in multiple cell lines and patient derived biopsy samples using western blotting, qRT-PCR and IHC. Transcript and protein expression values were correlated. CRISPR/Cas9-based knock out of the identified important protein in cisplatin-resistant cells restored drug induced phenotype. The patient derived cell (PDC) xenograft experiment suggests that knock out of the dysregulated protein induces cisplatin-mediated cell death and facilitate significant reduction of tumor burden. Mechanistically, the deregulated molecule regulates hippo signaling and activates YAP-1 target genes, which confers chemoresistance in OSCC. Following the discovery and validation of the protein target, we identified that the Radezolid (oxazolidinone group of antibiotics) represses the expression of the target protein and reverses drug resistance in OSCC-chemoresistant cell lines. The identified dysregulated molecule could be useful as a putative cancer marker explaining cisplatin-resistant development in OSCC cells.

## Materials and methods

### Ethics statement

This study was approved by the Institute review Board and Human Ethics committees (HEC) of Institute of Life Sciences, Bhubaneswar (84/HEC/18) and All India Institute of Medical Sciences (AIIMS), Bhubaneswar (T/EMF/Surg.Onco/19/03). The animal related experiments were performed in accordance to the protocol approved by Institutional Animal Ethics Committee of Institute of Life Sciences, Bhubaneswar (ILS/IAEC-147-AH/FEB-19).

### Cell culture and establishment of chemoresistant OSCC cells

The human tongue OSCC cell lines (H357, SCC-9 and SCC-4) were obtained from Sigma Aldrich, sourced from European collection of authenticated cell culture. PCR fingerprinting to establish the cell line authentication were done by the provider. All OSCC cell lines were cultured and maintained as described earlier [6].

### Generation of early and late cisplatin resistance cell lines

To establish a chemoresistant cell model system, OSCC cell lines (H357, SCC-9 and SCC-4) were initially treated with cisplatin at 1μM (lower dose) for a week and then the cisplatin concentration was increased gradually up to 15 μM (IC50 value) in a span of 3 months and further grown the cells at IC50 concentration until 8 month. Here, drugs efficiently eliminated the rapidly dividing cancer cells by inducing cell death, but poorly targeted the slowly dividing cells. Gradually, the poorly sensitive cells regained the normal growth cycle. Cells at the starting time were grouped as sensitive (CisS) and at 4 and 8 months of treatment were termed as early (Cis EarlyR) and late resistant (CisR Late R) cells respectively (Supplementary figure 1A).

### iTRAQ based proteomics analysis

Harvested cells (5×10^6^), from three time points (0M, 4M and 8M) were treated with RIPA buffer (Thermo Fisher Scientific, Cat #88665) supplemented with protease (details) and phosphatase inhibitor (Sigma, Cat # P0044,). Extracted cellular proteins from all three time points with appropriate technical and biological replicates were used in an isobaric tag for relative and absolute quantification (iTRAQ) experiment (**Fig. 1A & 1B**). Equal amount of proteins (100 μg) from all samples were taken for tryptic protein preparation following manufacturer’s instructions (AB Sciex, USA). Study samples with the tag details used for labelling in iTRAQ experiment are presented in (**Fig. 1B)**. Protein lysates were dried and dissolved using dissolution buffer and denaturant (2% SDS) supplied in the kit. Before trypsinization, proteins were reduced using tris-(2-carboxyethyl) phosphine (TCEP, 50 mM) at 60 °C for 1 hr, and cysteinyl residues were blocked using methyl methanethiosulfonate (MMTS, 200 mM). Trypsin treatment was performed using trypsin supplied by the manufacturer and incubating at 37°C for 16-20 hrs. Tryptic peptides were dried at 40°C using SpeedVac (LabConco, USA). Tagged tryptic peptides (∼250 μg) were subjected to strong cation exchange fractionation using a hand-held ICAT® Cartridge-cation-exchange system (Applied Biosystems, USA).

**Figure 1:**
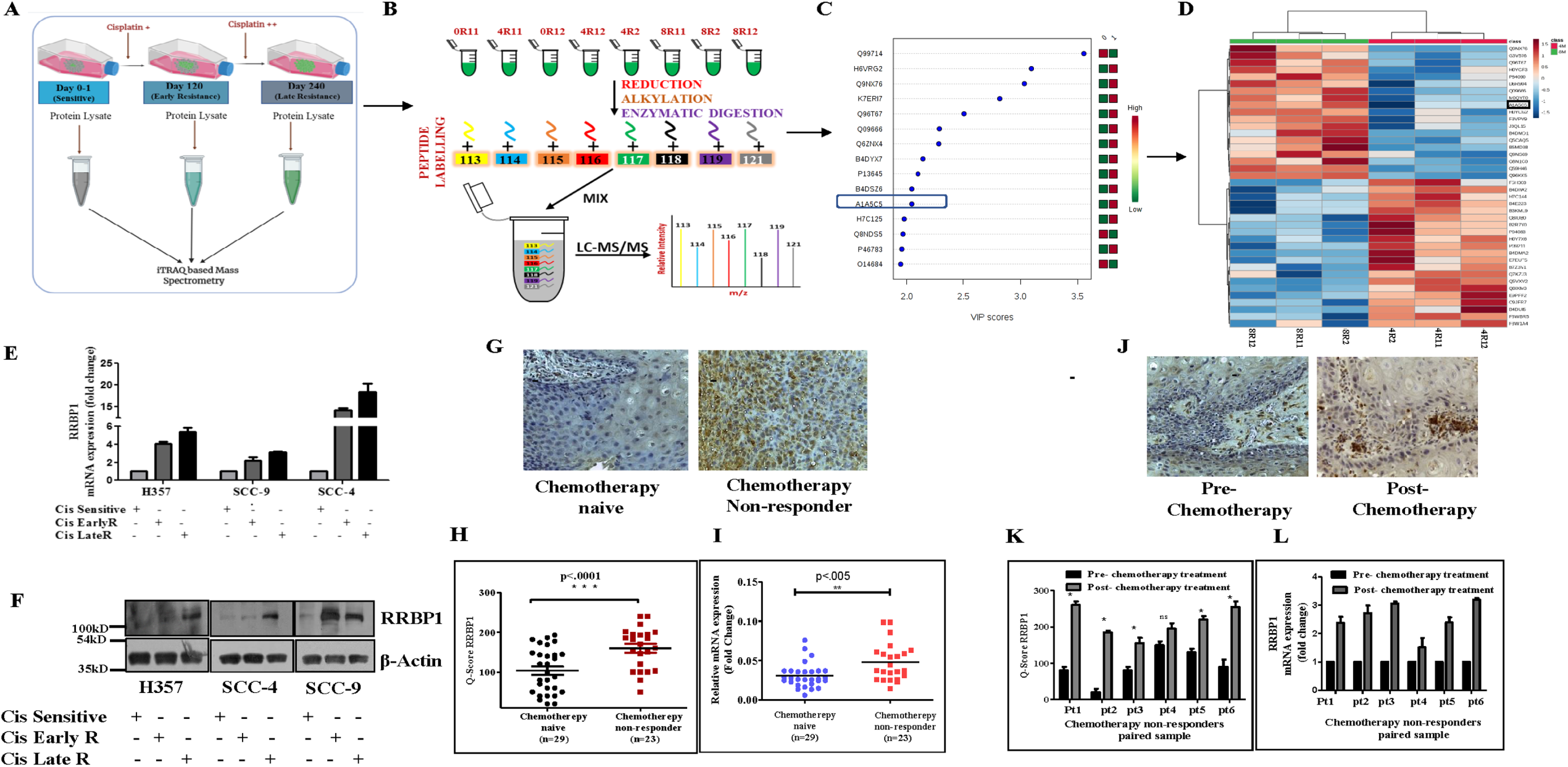
Global proteomics data revealed RRBP1 is upregulated in OSCC chemoresistant cells. **A)** Schematic representation of sensitive, early and late cisplatin resistant OSCC line for global proteomic profiling. The establishment of sensitive, early and late resistant cells is described in the materials and method section. **B)** The lysates were isolated from parental sensitive (H3457CisS), early (H357Cis Early R) and late (H357Cis Late R) cisplatin resistant cells and subjected to global proteomic profiling. The schematic diagram depicts the iTRAQ labeling strategy for proteomic analysis. 0R11 and 0R12 are technical replicates of H357CisS group, 4R11: 4R12 are technical and 4R2 is biological replicates of H357Cis EarlyR group, 8R11: 8R12 are technical and 8R2 is biological replicates of H357Cis LateR group. **C)** VIP score analysis of global proteomic profiling of sensitive, early (EarlyR) and late resistant cells (LateR). The uniport ID (A1A5C5) represents for human Ribosome Binding Protein. **D)** The dendrogram represents the deregulated genes from proteomic analysis (from early to late resistance normalized with sensitive). **E)** Relative mRNA (fold change) expression of RRBP1 was analyzed by qRT PCR in indicated acquired chemoresistant OSCC cells as compared to the sensitive counterpart (mean ±SEM, n=2). GAPDH was used as an internal control. **F)** Cell lysates from indicated resistant and sensitive OSCC cells were isolated and subjected to immunoblotting against RRBP1 and β-actin antibodies. **G)** Protein expression of RRBP1 was analyzed by immunohistochemistry (IHC) in chemotherapy-naïve and chemotherapy-non-responder OSCC tumors. **H)** Representative IHC Scoring for RRBP1 from panel G (Q Score =Staining Intensity × % of Staining), (Median, n=29 for chemotherapy-naïve and n=23 for chemotherapy-non-responder) *: P < 0.05. **I)** Relative mRNA expression of RRBP1 was analyzed by qRT PCR in different chemotherapy-non-responder OSCC tumors as compared to chemotherapy-naïve tumors (Median, n=29 for chemotherapy-naïve and n=23 for chemotherapy-non-responder). *: P < 0.05. **J)** Protein expression of RRBP1 was analyzed by immunohistochemistry (IHC) in pre- and post-TPF treated paired tumor samples for chemotherapy-non-responder patients. **K)** IHC Scoring for RRBP1 from panel J (Q Score =Staining Intensity × % of Staining), n=6. **L)** Relative mRNA expression of RRBP1 was analyzed by qRT PCR in pre- and post-TPF treated paired tumor samples for chemotherapy-non-responder patients (n=6). Pt represents each patient.

Each SCX fraction was resuspended in 20 μl of buffer (water with 0.1% formic acid) and introduced to easy-nanoLC 1000 HPLC system (Thermo Fisher Scientific, Waltham, MA) connected to hybrid Orbitrap Velos Pro Mass Spectrometer (Thermo Fisher Scientific, Waltham, MA). The nano-LC system contains the Acclaim PepMap100 C18 column (75 µm × 2 cm) packed with 3 μm C18 resin connected to Acclaim PepMap100 C18 column (50 µm × 15 cm) packed with 2 μm C18 beads. A 120 min gradient of 5% to 90% buffer B (0.1% formic acid in 95% Acetonitrile) and Buffer A (0.1% formic acid in 5% Acetonitrile) was applied for separation of the peptide with a flow-rate of 300 nl/min. The eluted peptides were electrosprayed with a spray voltage of 1.5 kV in positive ion mode. Mass spectrometry data acquisition was carried out using a data-dependent mode to switch between MS1 and MS2.

### Protein Identification and iTRAQ Quantitation

Protein identification and quantification was carried out using SEQUEST search algorithm of Proteome Discoverer Software 1.4 (Thermo Fisher, Waltham, MA, USA). Each MS/MS spectrum was searched against a human proteome database (UniProt, 89,796 total proteins, downloaded in April 2017). Precursor ion mass tolerance (20 ppm), fragmented ion mass tolerance (0.1 Da), missed cleavages (<2) for trypsin specificity, Carbamidomethyl (C), Deamidation (N and Q), Oxidation (M) and 8-plex iTRAQ label (N terminus and K) were set as variable modifications. The false discovery rate (FDR) at both protein and peptide level was calculated at 5%. The identified protein list with fold change values were exported to Microsoft Excel for further statistical analysis. Identified proteins from study samples and relative fold change values were selected for principal component analysis and a partial least square discriminate analysis model was built using MetaboAnalyst 3.0. All the mass spectrometry data files (.raw and .mgf) with result files were deposited in the ProteomeXchange Consortium (PXD0016977).

### Lentivirus production and generation of stable Cas9 over expressing chemoresistant lines

For Cas9 lentivirus generation, we transfected lentiCas9-Blast (Addgene, Cat#52962) along with its packaging psPAX2 and envelop pMD2G in HEK293T cells as lentiviral particles were generated by protocol described in Shriwas et al [7]. Chemoresistant cells were infected with lento Cas9 particles and treated with blastidicines hydrochloride (5 µg/ml, MP biomedical, Cat# 2150477). Single clones were selected, and Cas9 over expression was confirmed by Western blot using anti-Cas9 antibody (CST Cat #14697) (Supplementary Fig 3A). The lentiCas9-Blast vector was kindly deposited to Addgene by Feng Zhang lab [8].

### Generation of CRISPR based RRBP1 KO cell line

For generation of RRBP1 knockout cells, lentiviral vector expressing RRBP1 sgRNA (GGCGTTTCAGAATCGCCACA) was procured from Addgene (Cat# 92157). This lentiGuide-RRBP1-2 vector was kindly deposited by Alice Ting Lab [9]. Lentiviral particles were generated as described above using HEK293T cells. Stable clones of Cas9 overexpressing (Supplementary Fig 3A) chemoresistant cells (H357 CisR, SCC-9 CisR and patient derived cells PDC1) were infected with lentiGuide-RRBP1-2 for 48h in presence of polybrene (8 µg/ml), after which cells were treated with puromycin (2 μg/ml, Invitrogen, USA, Cat #A11138-03) for 7 days. Single clones were selected, and RRBP1 knockout was confirmed by Western blot using anti-RRBP1 antibody (Abcam, USA, Cat # ab95983). The RRBP1 KO clones were confirmed by cleavage detection assay (Supplementary Fig. 3B&C). The genomic cleavage efficiency was measured by the GeneArt® Genomic Cleavage Detection kit (Thermo Fisher Scientific, Cat # A24372) according to manufacturer’s protocol. Oligos used for this study are mentioned in supplementary table 3).

### Transient transfection with plasmids

For transient expression, H357 and SCC-9 cells were transfected with RRBP1 overexpression plasmids pcDNA4 HisMax-V5-GFP-RRBP1(Addgene:Cat#92150) using the ViaFect transfection reagent (Promega Cat# E4982). pcDNA4 HisMax-V5-GFP-RRBP1 was kindly deposited by Alice Ting Lab [9]. The cells were transfected for 48h, after which they were treated with cisplatin (5μM) followed by flow cytometry analysis (Annexin V PE/7-AAD Assay) and immunoblotting with anti-PARP and Anti β-actin. The transfection efficiency was determined by immunoblotting with Anti GFP (Abcam, USA, Cat # ab6556).

### OSCC patient sample

Biopsy samples of chemotherapy-naive patients (n=29, (OSCC patients that were never treated with any chemotherapy) and OSCC chemotherapy non-responders (n=23, OSCC patients were treated with neoadjuvant chemotherapy but never responded or partially responded) were collected from clinical sites. Study subject details with treatment modalities are presented in Supplementary Tables 1 and 2. Primary patient-derived cells (PDC) were isolated from harvested tissues of patients not responding to treatment and cultured [10].

### Organoid culture

Following earlier published methods with minor modification, chemoresistant lines (H357CisR, SCC-9 CisR) and patient derived cells (PDC1) were used for developing 3D organoid [11]. Organoid formation rate was defined as the average number of 50-mm spherical structures at day 14 that was divided by the total number of seeded cells in each well at day 0. During this experiment, at day 8 from establishing organoid culture, cisplatin (10μM) or DMSO (vehicle control) was used for treatment.

### Immunoblotting

Cell lysates were used for immunoblotting experiments as described earlier [12]. For this study, we used antibody against Anti β-actin (Abcam, USA, Cat#A2066), Anti RRBP1 (Sigma, Cat# HPA011924, Abcam USA, Cat# ab95983), Anti YAP(CST, Cat # 14074), Anti YAP-1^s-397^ (CST, Cat # 13619S), Anti YAP-1^s-127^ (CST, Cat # 13008), Anti PARP (CST, Cat #9542L), Anti p^s-139^-H2AX (CST, Cat # 9718S), Anti GFP (Abcam, USA, Cat # ab6556), and YAP/TAZ Transcriptional Targets Antibody Sampler Kit (CST, Cat#56674), anti Cas9 (CST, Cat # 14697).

### Patient Derived Xenograft

BALB/C-nude mice (6-8 weeks, male, NCr-Foxn1nu athymic) were purchased from Vivo biotech Ltd (Secunderabad, India) and maintained under pathogen-free conditions in the animal house. The patient-derived cells (PDC1) established from chemo non-responders was used for xenograft model [10]. The patient (PDC#1) was treated with TP (50 mg carboplatin and 20 mg paclitaxel for 3 cycles) without having any response. For xenograft experiment, cells (one million) were suspended in phosphate-buffered solution-Matrigel (1:1, 100 μl) and transplanted into upper flank of mice. The PDC1RRBP1KO cells were injected in the left upper flank and PDC1 WT cells were injected in right upper flank of same mice. After tumor reached volume of 50 mm^3^, we randomly divided these mice into 2 groups to treat with vehicle or inject cisplatin (3 mg/Kg) intraperitonially twice a week.

### RT-PCR and Real Time Quantitative PCR

Total RNA was isolated using RNA mini kit (Himedia, Cat# MB602) as per manufacturer’s instruction and quantified by Nanodrop. RNA (300 ng) was used for reverse transcription by using first cDNA synthesis (VERSO CDNA KIT Thermo Fisher Scientific, Cat # AB1453A) and qRT-PCR was carried out using SYBR Green master mix (Thermo Fisher scientific Cat # 4367659). GAPDH was used as a loading control and complete primer details used in this article are listed in supplementary table 3.

### Immunohistochemistry

OSCC patients tumor and mice tumors were isolated for paraffin embedding for immunohistochemistry following previously described method [7]. Antibodies for RRBP1 (Sigma, cat# HPA011924), Ki67 (Vector, Cat# VP-RM04), cleaved caspase (CST, cat # 9661S), CTGF (Santa Cruz Cat# SC 25440), Survivin (CST Cat# 2808) were used for immunohistochemistry. Q-score was determined by protocol as described in Maji et al [10].

### Annexin V PE/7-ADD Assay

Apoptosis and cell death were measured by staining with Annexin V Apoptosis Detection Kit PE (eBioscience™, USA, Cat # 88-8102-74) as described earlier [10].

### Immunofluorescence

Cell (10^3^) were seeded on coverslip and allowed to adhere to the surface. The adhered cells were fixed in 4% formaldehyde for 15 min and permeabilize with 1X permeabilization buffer (eBioscience™, USA, Cat # 00-8333-56) followed by blocking with 3% BSA for 1h at room temperature. Then cells were incubated with primary antibody overnight at 4°C, washed thrice with PBST followed by 1h incubation with Alexa fluor conjugated secondary antibody (Thermo Fisher Scientific Cat # 11008, 21244) then again washed thrice with 1XPBST, after which coverslips were mounted with DAPI (Slow Fade ® GOLD Antifade, Thermo Fisher Scientific, Cat # S36938). Images were captured using a confocal microscopy (LEICA TCS-SP5).

### Colony formation assay

For colony forming assay, cells (500) were seeded in 6 well plate and treated with DMSO, Cisplatin (*cis*-diammineplatinum (II) dichloride, Sigma, Cat#479306), Radezolid (MedChemExpress USA, Cat #HY-14800) or in combination then allowed for 10 days to grow. The colonies were stained with 0.5% crystal violet and counted by ImageJ software.

### Validation of relative expression levels of YAP target gene with RRBP1

The relative expression levels of YAP target gene with RRBP1 in HNSCC patient tumors were validated using GEPIA (http://gepia.cancer-pku.cn/detail.php?gene=RRBP1), online analysis software based on the TCGA database and Genotype, using |log_2_FC|≥1 and P≤0.05 as the cut-off criteria.

### Insilco molecular docking

The Radezolid structure was obtained from DrugBank (https://www.drugbank.ca/). The ligand was prepared by LigPrep module in Schrodinger molecular modeling software. Since there is no crystal structure available for the target protein RRBP1 in Protein Databank (PDB), a homology model of RRBP1 was built using Modller9.22 software. The protein template for homology modeling was selected using DELTA-BLAST. Two templates (PDB ID 6FSA, and 5TBY) were selected and multiple sequence alignment with RRBP1 sequence was done for the modeling. Ten models were built using Modeller9.22 and the best model was chosen on MolPDF score. The selected model was imported in the Maestro module of Schrodinger software to prepare it for docking. The hydrogens were added, bond orders and ionization states were assigned and charges were calculated for the atoms of the RRBP1 protein. The active site of RRBP1 protein was identified by SiteMap and a cavity with 772.7 A3 was selected for docking. The docking study was done using the Glide module with extra precision (XP) mode.

### Statistical analysis

All data points are presented as mean and standard deviation and Graph Pad Prism 5.0 was used for calculation. The statistical significance was calculated by one-way variance (one-way ANOVA), Two-Way ANOVA and considered significance at P≤0.05.

## Results

### Establishment of chemoresistance OSCC cells

To identify the key resistance triggering molecules, we have established cisplatin resistant cells by prolonged treatment of cisplatin to OSCC cell lines as described in the method section. Monitoring cisplatin-induced cell death in three stages (CisS, Cis EarlyR and Cis LateR) of H357, SCC-9 and SCC-4 cells by flow cytometry assay showed Cis LateR achieved complete acquired resistance and Cis EarlyR achieved partial resistance (Supplementary Fig 1A-B)

### RRBP1 expression is elevated in chemoresistant OSCC

From the adopted global proteomics experiment including H357 CisS, H357 Cis EarlyR and H357 Cis LateR, a set of **247 proteins** were identified and **44** showed dysregulations (log_2_(resistance/sensitive)>±1.0 and VIP score>1.6) (Fig. 1 A-B and supplementary table 4). Principal Component Analysis reveals that (PCA), all the identified proteins as variables with their fold change values are grouped into two different cluster (Supplementary Fig.2A). The variable importance in the projection (VIP) values were also applied to identify deregulated proteins with cut-off value >2 (Fig. 1C). From literature mining, we selected one of these important deregulated molecules i.e. Ribosome Binding Protein 1 for further validation. The dendrogram indicates that RRBP1 expression is elevated during the development of cisplatin resistance (Fig 1D). Further we checked and confirmed the fragmentation spectra of RRBP1 protein quantified by peptide ALNQATSQVES (Supplementary Fig 2B-C). Based on these observations, we went for further validation of RRBP1 and to elucidate its potential role in acquired cisplatin resistance. In independent sample sets, we monitored the RRBP1 expression in H357, SCC-9 and SCC-4 OSCC tongue cell lines showing different acquired chemoresistant patterns (H357 Cis EarlyR, H357 Cis LateR, SCC-4 Cis EarlyR, SCC-4 Cis LateR, SCC-9 Cis EarlyR and SCC-9 Cis LateR, R indicates Resistance). Expressions of RRBP1 at protein and mRNA levels were found to be up-regulated in Cis EarlyR and Cis LateR cells with respect to CisS (cisplatin sensitive) counterparts in all cell lines (Figure 1E-F). A similar profile of RRBP1 was also observed in tumor isolated from drug naïve (freshly diagnosed OSCC tumors) and patients not responding or partially responding to neoadjuvant chemotherapy (TPF) (Figure 1G-I). In drug-naive and post-treated paired tumor samples not responding to neoadjuvant chemotherapy treatment, we observed higher expression of RRBP1 in post chemotherapy treated tumors (Figure 1J-L). From the in-vitro and tumor samples isolated from OSCC patients presenting with chemoresistance, consistently higher RRBP1 abundance was observed.

### RRBP1 dependency in chemo-resistant cell lines and patient-derived cells (PDC)

We generated RRBP1 knock out clones in H357 CisR and SCC-9 CisR which was confirmed by cleavage detection assay (Supplementary Fig 3 A-C). The RRBP1 KO chemoresistant cells restored cisplatin-mediated cell death in H357 CisR and SCC-9 CisR (Figure 2A, B) After knocking out RRBP1 from patient-derived tumor primary cells not responding to TP, PDC1 cells reversed resistance and became sensitive to cisplatin-induced cell death (Figure 2C). In addition to this, transient over expression of RRBP1 in H357CisS and SCC-9 CisS cells using over-expression construct (pcDNA4 HisMax-V5-GFP-RRBP1), showed the development of cisplatin resistance (Fig. 3A-F). This observation indicated an RRBP1 dependency of OSCC chemoresistant cells.

**Figure 2:**
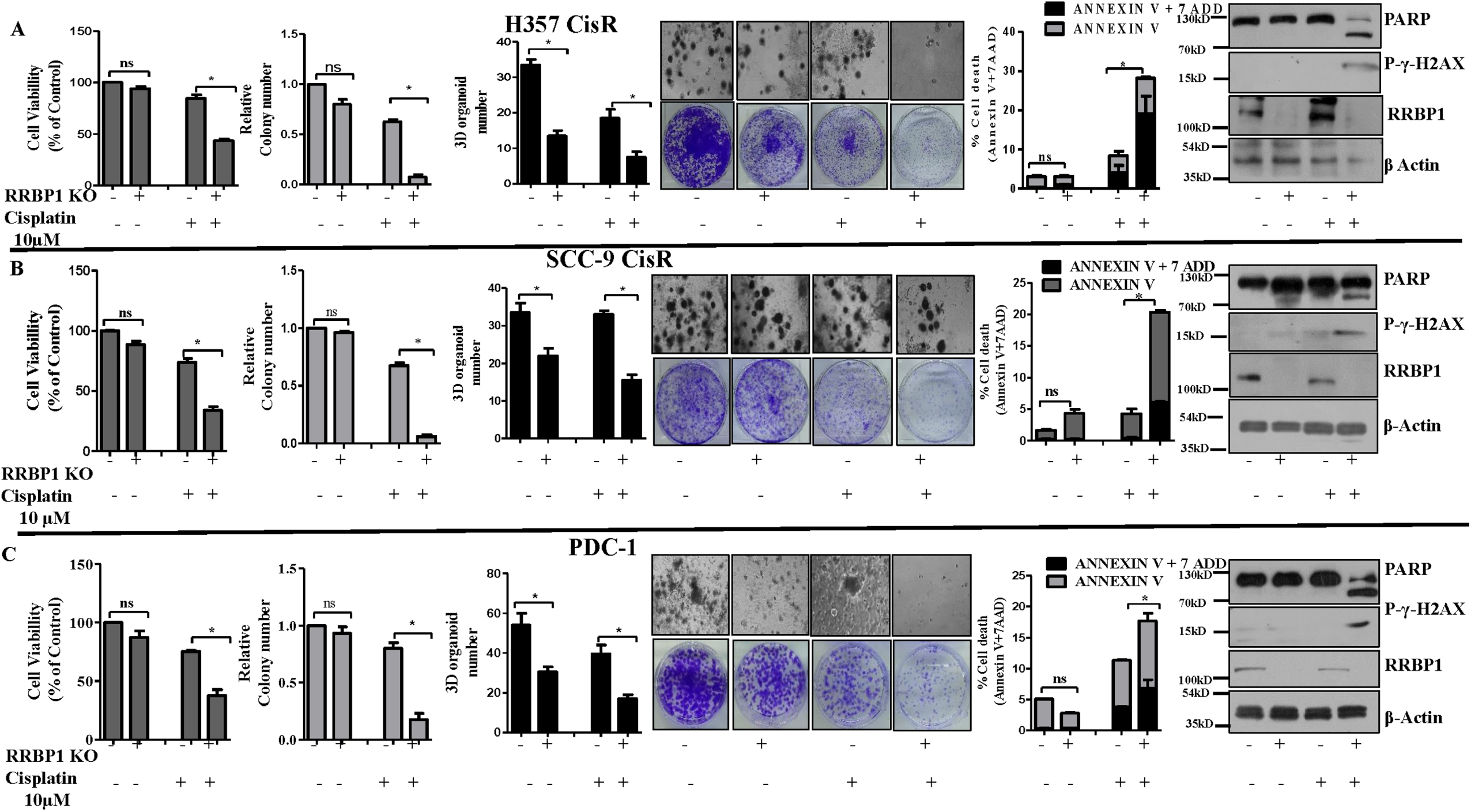
RRBP1 knock out sensitized chemoresistant resistant cells to cisplatin. **A)** RRBP1 KO cells in human OSCC line H357CisR were generated using a lentiviral approach as described in materials and methods. **1**^**st**^ **panel**: RRBP1KO and RRBP1WT cells were treated with cisplatin (10μM) for 48 hours and cell viability was determined using MTT assay (Mean ±SEM, n=3) *: P < 0.05. **2**^**nd**^ **panel**: RRBP1KO and RRBP1WT cells were treated with cisplatin (10μM) and anchorage dependent colony forming assay was performed as described in materials methods. The bar describes the relative colony number in each treatment group (Mean ±SEM, n=3) *: P < 0.05. **3**^**rd**^ **panel**: RBP1KO and WT cells were treated with cisplatin (10μM) and 3D organoid assay was performed as described in materials and methods. The bar diagram represents the number of organoids in each treatment group (Mean ±SEM, n=3) *: P < 0.05. **4**^**th**^ **panel:** representative images of anchorage dependent colony forming assay (lower panel) and 3D organoid assay (upper panel) as described in 2^nd^ and 3^rd^ panel. **5**^**th**^ **panel**: RRBP1KO and WT cells were treated with cisplatin (10μM) for 48h, after which cell death was determined by annexin V/7AAD assay using flow cytometer. Bar diagrams indicate the percentage of cell death with respective treated groups (Mean ±SEM, n=2). **6**^**th**^ **panel:** RRBP1KO and WT cells were treated with cisplatin (10μM) for 48h and immunoblotting was performed with indicated antibodies. **B)** RRBP1 KO cells in human OSCC line SCC-9 were generated using a lentiviral approach as described in materials and methods. Similar experiments were conducted with SCC-9 CisR RRBP1 KO and SCC-9 CisR RRBP1WT as described in all panels of section A. **C)** RRBP1 KO cells in PDC1 (patient derived cells) were generated using a lentiviral approach as described in materials and methods. Similar experiments were conducted with PDC1 RRBP1 KO PDC1 RRBP1WT as described in all panels of section A.

**Figure 3:**
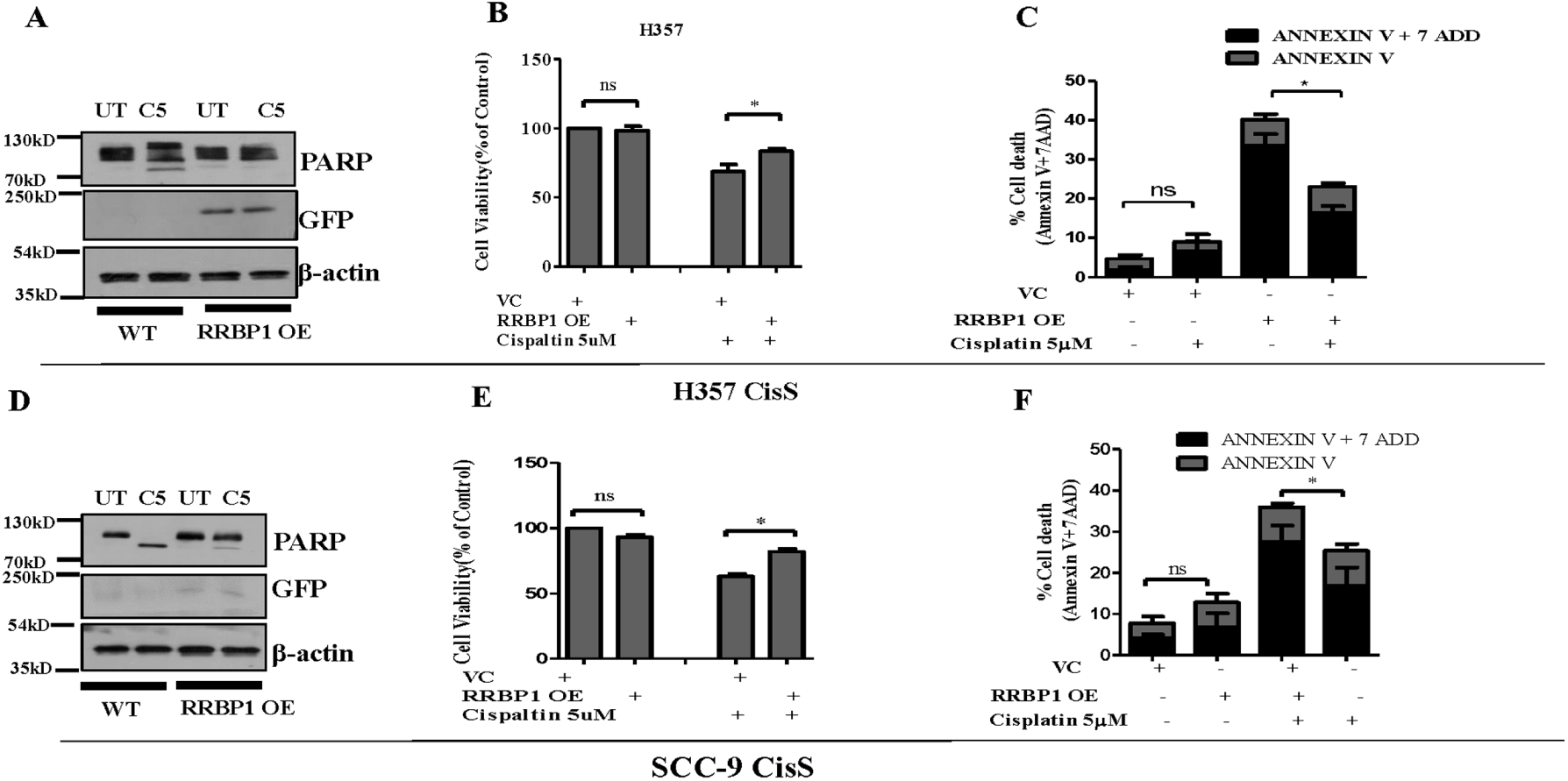
Overexpression of RRBP1 in human OSCC lines results in development of cisplatin resistance. **A)** H357CisS were transfected with RRBP1 overexpression vector (pcDNA4 HisMax-V5-GFP-RRBP1) and treated with cisplatin (5 μM) for 48h, after which immunoblotting was performed against indicated antibodies. GFP expression indicates the transfection efficiency of RRBP1 overexpression construct. **B)** Cells were treated as described in A panel and cell viability was determined by MTT assay (Mean ±SEM, n=3) *: P < 0.05. **C)** Cells were treated as described in A panel and cell death was determined by annexin V/7AAD assay using flow cytometer. Bar diagrams indicate the percentage of cell death with respective treated groups (Mean ±SEM, n=3). **D-F)** SCC-9 CisS cells were transfected with RRBP1 overexpression vector (pcDNA4 HisMax-V5-GFP-RRBP1) and treated with cisplatin (5μM) for 48h and experiments were performed as described in panel A-C.

### RRBP1 regulates YAP-1 expression in chemoresistant cells

The deregulated proteins, identified from global proteomics analysis, were converted to gene list and a functional analysis was carried out using Ingenuity Pathway Analysis (IPA). IPA analysis of functional pathways in acquired chemo-resistance cells showed highest down-regulation of Hippo signaling (Supplementary Fig. 4A-B). Hippo pathway negatively regulates the activity of its downstream transcriptional co-activators, Yes-associated protein 1 (YAP-1) and Transcriptional coactivator with PDZ-binding motif (TAZ). The active YAP-1/TAZ translocate to the nucleus and binds with TEA domain family member (TEAD), which in turn transcribes genes that promotes cell proliferation and inhibit apoptosis. Dysregulated Hippo signaling promotes malignancy in cancer cells and mediate chemoresistance in different neoplasms.

Expression of YAP-1 and its target genes (CYR61, CTGF, Jagged 1, AXL, Integrin β2, IGFBP3 and Laminin B2) monitored in RRBP1 KO chemoresistant cells and WT, were found to be significantly downregulated both at mRNA and protein levels (Figure 4A and 4B) in KO cells. Moreover, our confocal microscopy data suggest that nuclear YAP-1 expression is significantly reduced in RRBP1 KO cells as compared to wild type cells (Fig.4C). Hippo signaling is tightly regulated by MST1/2 and LATS1/2, which phosphorylates the YAP-1 at Ser127 (Hippo on) resulting in its cytoplasmic retention and proteasome-mediated degradation. Similarly, phosphorylation at Ser397 of YAP-1 by LATS1/2 creates a phospho□degron motif for β□TrCP binding followed by proteasomal degradation [13]. We did not find any increase in phosphorylated YAP-1, rather, lower expression of p-YAP (at Ser-127 and Ser-397) were observed in RRBP1 KO as compared to WT cells (Figure 4D). YAP-1 m-RNA expression in WT and RRBP1 KO cells were found to be similar (Figure 4E). Further, association analysis of RRBP1 m-RNA levels with YAP-1 target gene (CTGF, CYR61, Jagged 1, AXL, TEAD1, COL1A1,DCN, NUAK1,FOXO1, AMOTL2, PCDH7 and MYOF) in the TCGA (The Cancer Genome Atlas) HNSCC cohort using GEPIA showed a positive correlation (r≥0.3) (Fig. 4F). Collectively these data demonstrated that loss of RRBP1 impair YAP-TEAD target gene expression, henceforth RRBP1 regulate YAP1 expression.

**Figure 4:**
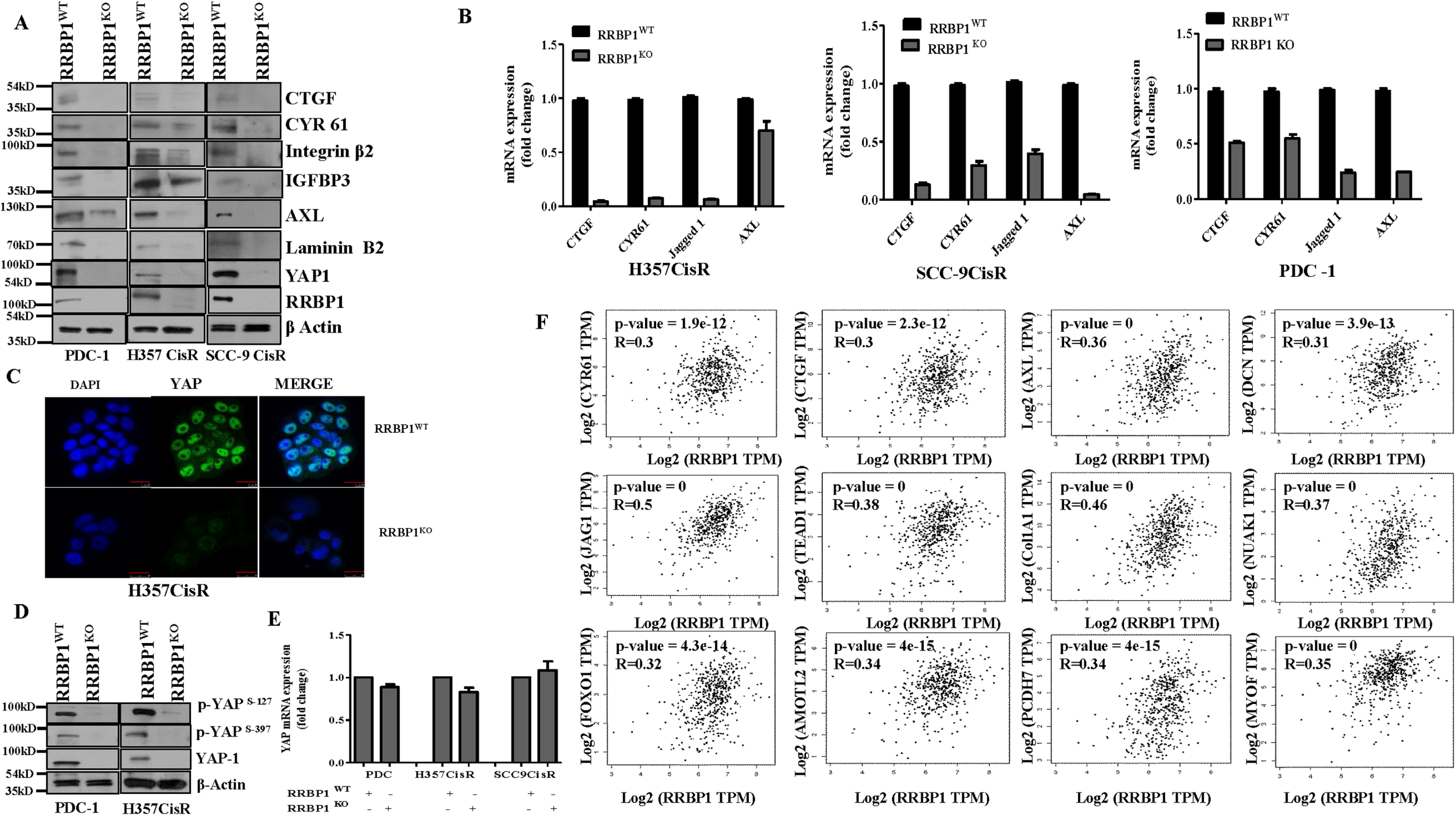
RRBP1 regulates YAP in chemoresistant OSCC. **A)** Cell lysates from indicated RRBP1KO and WT cells were isolated and subjected to immunoblotting against indicated antibodies. **B)** Relative mRNA (fold change) expression of indicated genes was analyzed by qRT PCR in indicated RRBP1 KO and RRBP1WT chemoresistant cells (mean ±SEM, n=3). GAPDH was used as an internal control. **C)** H357CiSR RRBP1KO and H357 CisR RRBP1WT cells were cultured and immunostaining were performed using the anti-YAP-1 antibody as described in materials and methods. Images were acquired using confocal microscopy (LEICA TCS-SP8). **D)** Cell lysates from indicated RRBP1KO and RRBP1WT were isolated and subjected to immunoblotting against indicated antibodies. **E)** Relative mRNA (fold change) expression of RRBP1 was analyzed by qRT PCR in indicated cell lines with RRBP1 KO and RRBP1 WT (mean ±SEM, n=3). **F)** Expression correlation of RRBP1 and YAP-1 target genes (mRNA) analyzed in the HNSCC TCGA cohort. (CTGF, CYR61, Jagged 1, AXL, TEAD1, COL1A1, DCN, NUAK1, FOXO1, AMOTL2, PCDH7 and MYOF). Correlation was analyzed using Spearman’s correlation coefficient test, n = 520. The analysis was performed in Gene Expression Profiling Interactive Analysis (GEPIA) platform.

### Knock out of RRBP1 significantly induced cisplatin-mediated apoptosis in chemoresistant patient-derived xenograft

To evaluate in-vivo efficacy of knocking out RRBP1 in restoring cisplatin-induced cell death in chemoresistant OSCC, we established xenograft tumors in nude mice using PDC isolated from tumors of a chemotherapy-non-responder patient(Fig 5A) (patient# 1, table 1). The PDC1 WT cells were implanted in the right flank of athymic nude mice and PDC1RRBP1KO cells were implanted in the left flank of same mice. We observed a reduction in tumor growth and size in RRBP1 KO group as compared to WT PDC1. Treating with cisplatin (3 mg/kg) significantly reduced the tumour burden in case of RRBP1 KO group (Figure 5 B-D). Immunohistochemistry analysis of harvested tumors showed significantly higher apoptosis levels, reduced expression of YAP-1 and its target genes in RBBP1KO groups treated with cisplatin (Figure 5E).

**Figure 5:**
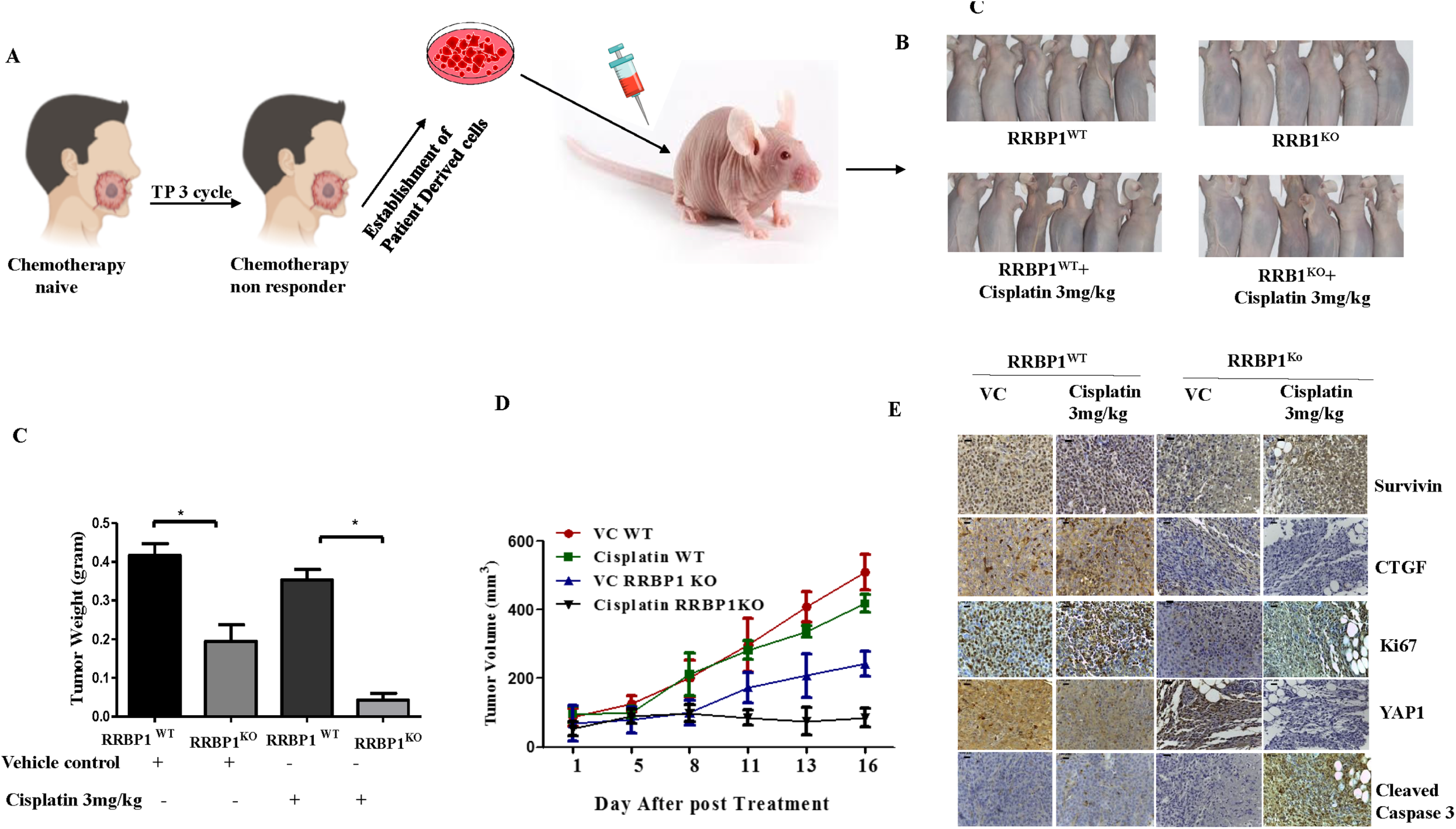
Knock out of RRBP1 restores cisplatin induced cell death in chemoresistant xenografts. **A)** Schematic representation of establishment of PDX, using patient derived cells isolated from chemotherapy-non-responder patient **B)**. RRBP1 WT cells were injected to the right flank of athymic nude mice and PDC1RRBP1KO cells were injected to the left flank of same mice as described in materials and methods. After which mice were treated with either vehicle control or 3 mg/Kg of cisplatin in two different groups (twice a week). At the end of the experiment’s mice were sacrificed and images were captured (n=6). **C)** At the end of experiments, tumor weight was measured and represented in bar diagram (mean ± SEM, **P < 0.05, n = 6). **D)** Tumor growth was measured in the indicated time point using digital slide caliper and plotted as a graph (mean ± SEM, n = 6). **E)** After completion of treatment, tumors from each group were fixed with formalin, and paraffin-embedded sections were prepared to perform immunohistochemistry with indicated antibodies.

### Radezolid, a potential inhibitor of RRBP1 restores cisplatin-induced cell death in chemoresistant OSCC

As potential inhibitors of RRBP1 are missing the in literature. From PubChem and drug bank database search, it seemed that Radezolid, a second generation oxazolidinone antibiotic could be used as a potential candidate to target RRBP1. The Insilco molecular docking study suggested that the molecule docks well at the active site (Fig 6 A-B) making multiple hydrogen bond interactions. The molecule shows a docking score of -8.0 indicating it is having a high affinity for the RRBP1 protein. There are three hydrogen bonds interacting with RRPB1 and Radezolid. The first hydrogen bond with GLU78 while the nearby amide makes a hydrogen bond with PRO115. The other amide makes a hydrogen bond interaction with ARG101 of RRBP1 (Fig 6 C-D). The docking score and three hydrogen bond interaction showed that Radezolid have potential binding affinity to inhibit the RRBP1 expression. Next, RRBP1 expression was monitored in chemoresistant cells (H357CisR, SCC-9 CisR and PDC1) treated with Radezolid. We observed a dose dependent (≥5 μM) lowering of RRBP1 protein expression with treatment of Radezolid (Fig. 6E). Interestingly, our qRT-PCR data suggests that Radezolid treatment did not affect the mRNA expression of RRBP1 in chemoresistant cells (Fig. 6F). However, Radezolid treatment significantly reduced the expression of YAP-1 target genes in H357CisR and SCC-9CisR cells (Fig. 6G). Further, we evaluated if treatment of Radezolid can overcome cisplatin resistance in OSCC. Our data suggest upon combination effect of Radezolid and Cisplatin treatment in cisplatin-resistant OSCC cells and PDC1, we observed a reversal of chemoresistance (Figure 7 A-C). Apoptosis induced by this combinatorial effect was confirmed from a significant increase in cleaved PARP, and increased p-γH2AX level. Our ongoing effort is to understand the in vivo efficacy of Radezolid in chemoresistant PDX.

**Figure 6:**
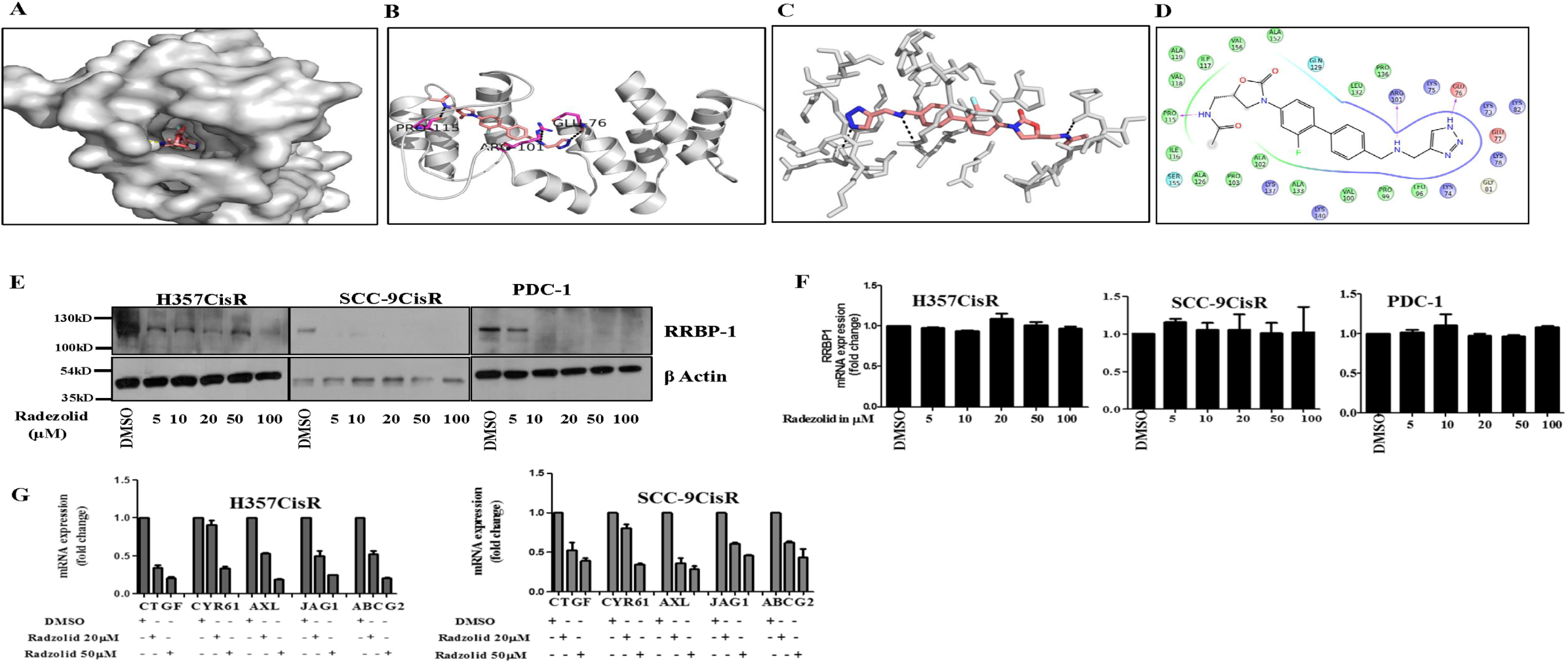
Radezolid represses RRBP1 protein expression and regulate YAP target gene in chemoresistant OSCC. **A)** Molecular docking of RRBP1 with Radezolid compounds carried out in Glide and representation in surface view receptor of drug interaction **B)** Ribbon model structure of RRBP1 showing the hydrogen bonding interaction with Radezolid at PRO-115, ARG-101 and GLU-76. Hydrogen bonds are shown as dashed lines. **C)** Active site residues within 5 Angstrom of RRBP1 and Radezolid interactions. **D)** Ligand plot showing interaction of the Radezolid interaction with different residues of RRBP1 within 5 angstrom distance. **E)** Indicated chemoresistant OSCC cells were treated with Radezolid in a dose dependent manner for 48h, after which lysates were isolated to perform immunoblotting against β-actin and RRBP1 **F)** Indicated cells were treated with Radezolid in a dose dependent manner for 48h and qRT-PCR was performed to determine the relative mRNA expression (fold change) of RRBP1. GAPDH was used as an internal control. **G)** Chemoresistant cells (H357CisR and SCC-9CisR were treated with indicated concentration of Radezolid and qRT-PCR was performed to determine the relative mRNA expression of YAP-1 indicated target genes. GAPDH was used as an internal control.

**Figure 7:**
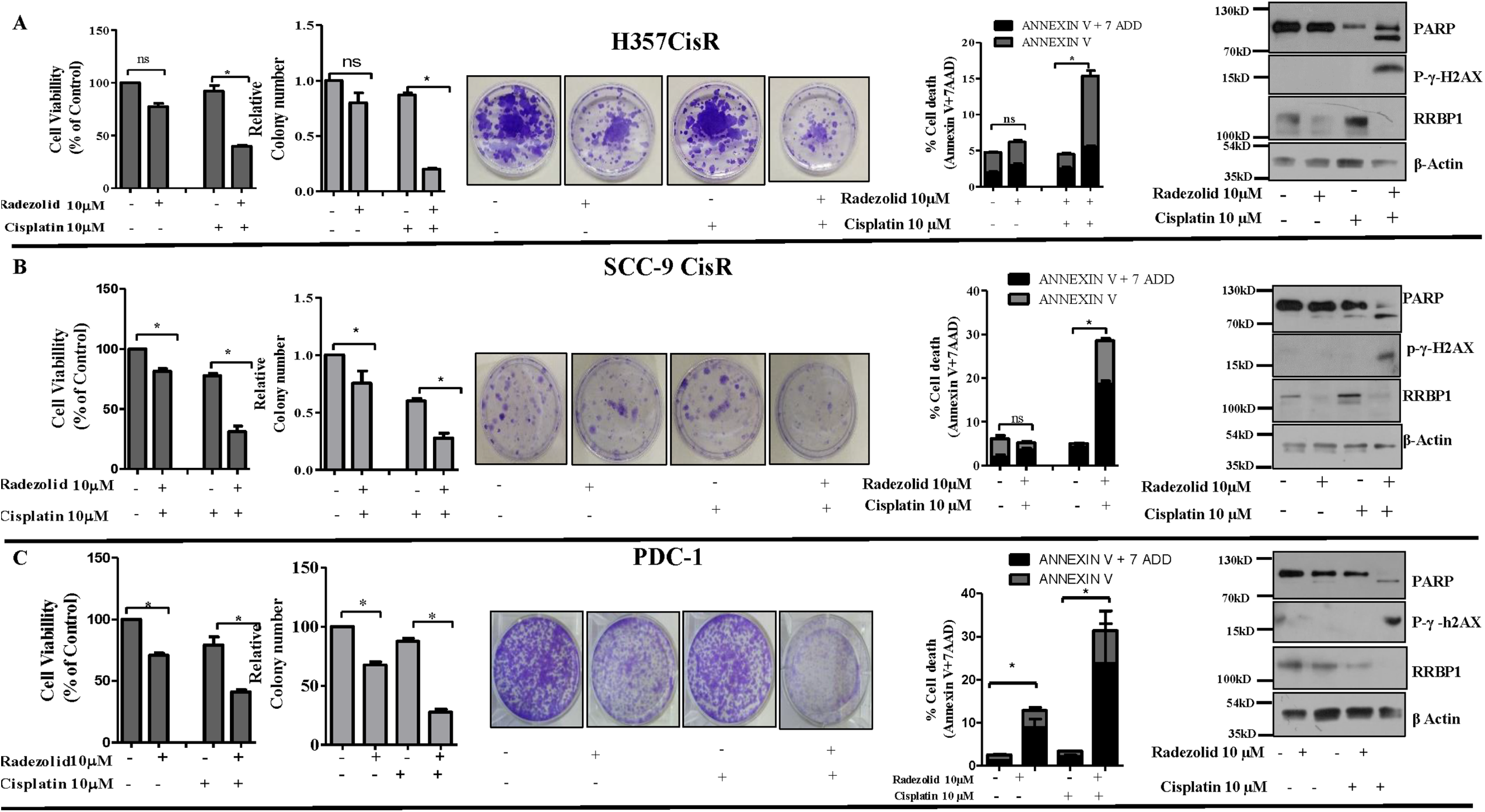
Radezolid (oxazolidinone group antibiotic) restored cisplatin-induced cell death in chemoresistant OSCC. **A). 1**^**st**^ panel: Cisplatin resistant OSCC line H357 CisR cells were treated with Radezolid (10µM) and cisplatin(10µM) alone or in combination for 48h and cell viability was determined using MTT assay (Mean ±SEM, n=3) *: P < 0.05. (A **2**^**nd**^ panel: H357 CisR cells were treated with Radezolid (10µM) and cisplatin(10µM) after which anchorage dependent colony forming assay was performed as described in materials methods. The bar describes the colony number as compared to vehicle treated H357CisR (cells (Mean ±SEM, n=3) *: P < 0.05. **3**^**rd**^ **panel**: Images of the anchorage dependent colony forming assay described in 2^nd^ (panel). **4**^**th**^ **panel**: H357 CisR cells were treated with Radezolid (10µM) and cisplatin(10µM) for 48h, after which cell death was determined by annexin V/7AAD assay using flow cytometer. Bar diagrams indicate the percentage of cell death from a panel with respective treated groups (Mean ±SEM, n=3). **5**^**th**^ **panel**: H357 CisR cells were treated with Radezolid (10µM) and cisplatin (10µM) for 48h, after which immunoblotting was performed with indicated antibodies. **B)** Cisplatin resistant OSCC line SCC-9 CisR cells were treated with Radezolid (10µM) and cisplatin(10µM) after which different assays were performed as described in section A. **C)** PDC-1 cells were treated with Radezolid (10µM) and cisplatin(10µM) after which different assays were performed as described in section A.

## Discussion

Earlier, cisplatin resistance models have been successfully established by prolonged treatment of drugs to cancer cell lines representing various neoplasms [14]. These models can be broadly divided into two groups, I) clinically relevant model, where cells were grown with lower doses of drug adopting a pulse treatment strategy [15, 16], II) High-level laboratory models, where cells are continually grown in presence of drugs with dosage escalation from lower dose to IC50 [17, 18]. The advantage of clinically relevant models is that it mimics the chemotherapy strategies in patients; on the other hand, its resistance pattern is very inconsistent. High-level laboratory models showed consistent resistant pattern and are generally preferred to study the mechanism of chemoresistance in cancer cells. All these studies engage the parental sensitive cells and late drug resistant cells to understand the molecular mechanism for chemoresistance. Here, to explore the causative factors of chemoresistance in OSCC, we have established and characterized sensitive, early and late cisplatin-resistant cells. Using global proteomic profiling of oral cancer cells with different grades of resistance to cisplatin, we have identified and validated that Ribosomal binding protein 1 (RRBP1) is one of the critical proteins responsible for resistance development. RRBP1 is localized in the rough endoplasmic reticulum (rER) and supposed to play a role in secretion of newly synthesized protein [19]. RRBP1 is reported to be over expressed in breast [20], lung [19], colorectal [21], esophageal [22], endometrial [23], prostate [24] and ovarian cancer [25] patient tissues. RRBP1 expression is associated with the disease progression and also envisaged as an unfavourable post-operative prognosis [21]. Here, we established that inhibiting RRBP1 expression has cisplatin resistance rewiring potential making it susceptible to cisplatin. To the best of our knowledge, this is the first study to demonstrate that elevated RRBP1 in cisplatin resistant OSCC cancer cells could be reversed/manipulated to make them susceptible to cisplatin treatment.

RRBP1 proteins, critical for translation, transportation and secretion of secretory proteins, anchors to the rough endoplasmic reticulum and also present in cytoplasm and nucleus [26, 27]. It is critical for translation, transportation and secretion of secretory proteins [27]. It plays an important role in augmenting collagen synthesis and secretion at the entry of secretory compartment. Knocking down RRBP1 in human fibroblast results in a significant reduction of secreted collagen. Electron microscopy study suggests lesser interaction of ER and ribosome in RRBP1 knocked down cells [28]. RRBP1 also mediates the targeting of certain m-RNAs to the ER. It was detected in mass spectrometry analysis of ER bound polysomes. The m-RNA targeted to ER generally has a signal receptor peptide. SRP of the ER destined m-RNA binds to RRBP1 (SRP receptor proteins on ER) through its single transmembrane domain and a large carboxyterminal lysine-rich domain. Interestingly, it was found that knocking down RRBP1 can inhibit the translation-dependent and translation independent ER association of specific mRNAs encoding calreticulin and alkaline phosphatase.

In this study, we observed a significant reduction of YAP-1 in RRBP1 knock out cells, limited or no YAP-1 phosphorylation, indicating that it is not degraded by proteasomal degradation. Our results indicate that RRBP1 might be playing a role in the translation of YAP-1 mRNA. In absence of RRBP1, the translation of YAP-1 is partially blocked (Fig. 8). It is well established that dysregulated Hippo signaling promotes malignancy in cancer cells [29, 30]. Recently, Hippo signaling has also been correlated to mediate chemoresistance in different neoplasms to several chemotherapeutic drugs [31]. Similarly, phosphorylation at S397 of YAP-1 by LATS1/2 creates a phospho□degron motif for β□TrCP binding followed by proteasomal degradation [13]. Overall, dephosphorylated YAP-1 (Hippo off) translocate to the nucleus to transcribe the YAP-1 target genes.

**Figure 8:**
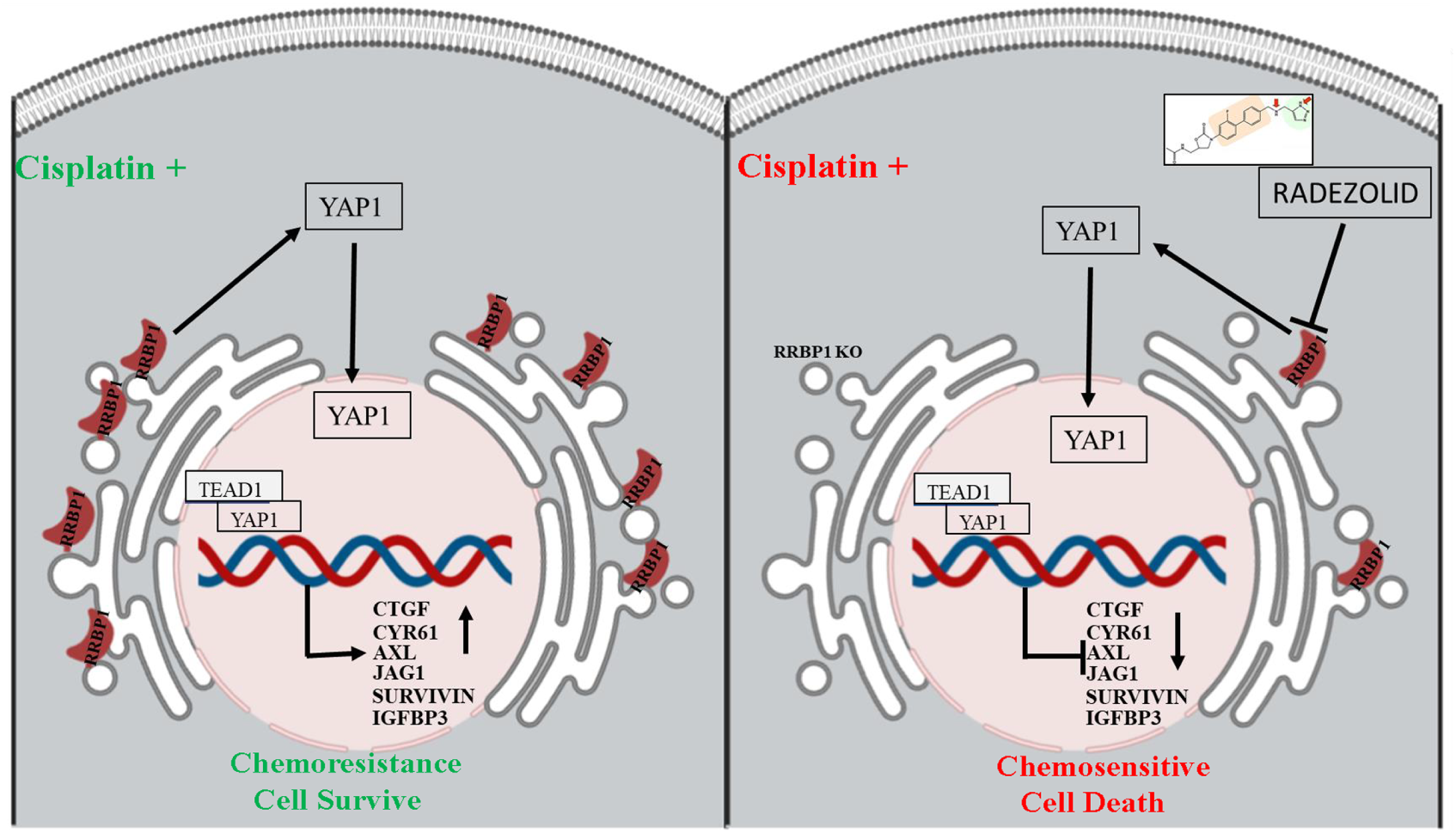
Schematic presentation of the mechanism by which RRBP1 mediates chemoresistance in OSCC. RRBP1 confers cisplatin resistance in OSCC via YAP-1 and its target gene. In presence of cisplatin RRBP1 expression elevated which activate YAP-1. As a result, cisplatin showing resistance and no cell death. Radezolid is an inhibitor of RRBP1 which repress RRBP-1 expression and induces cisplatin mediated cell death.

We additionally provide evidence that a second generation Oxazolidinones (Radezolid), synthetic antibiotic affecting the initiation phase of bacterial protein synthesis (under development by Melinta Therapeutics Inc) represses RRBP1 in chemoresistant cells. Further, preclinical studies are underway to determine the toxicity profile, pharmacokinetic properties of Radezolid. In recent future, we will evaluate the in-vivo efficacy of Radezolid, if it can reverse the cisplatin resistance in OSCC.

In conclusion, our findings suggest that blocking the RRBP1 expression in cisplatin resistant cells can be a viable strategy to overcome cisplatin resistance. We further demonstrated that Radezolid could be useful to reverse cisplatin resistance in acquired chemoresistant lines and PDX models. Further studies are warranted to establish the mechanism of action.

## Acknowledgements

Grant support: This work is supported by Institute of Life Sciences, Bhubaneswar intramural support and ICMR (5/13/9/2019-NCD-III) and. RD is thankful to Ramalingaswami Fellowship. SM is UGC-JRF, OPS is a UGC-SRF. Core support from International Centre for Genetic Engineering and Biotechnology is highly acknowledged.

