## Supplementary Figure 1 for "RRBP1 rewires cisplatin resistance in Oral Squamous Cell Carcinoma by regulating YAP-1"

### Supplementary Fig 1

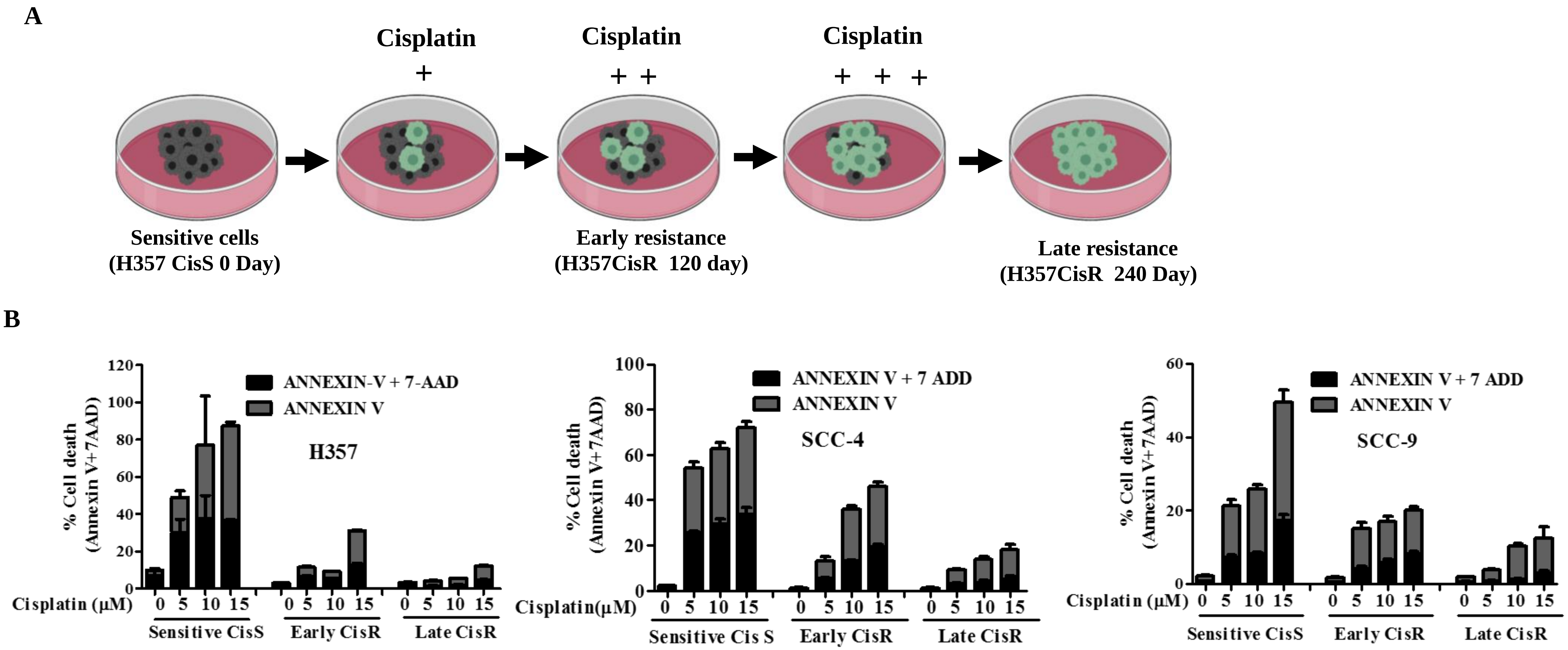


**Supplementary figure 1: Characterization of sensitive, early and late OSCC cisplatin resistant lines: A)** Schematic presentation of establishing sensitive, early and late cisplatin resistant cancer lines **B)** Sensitive, early and late cisplatin resistant pattern (CisS, CisR120 day and CisR 240 day) of H357, SCC-9 and SCC-4 cells were treated with indicated concentration of cisplatin for 48h, after which cell death was determined by annexin V/7AAD assay using flow cytometer. Bar diagrams indicate the percentage of cell death with respective treated groups (Mean ±SEM, n=3)
