## Supplementary Figure 2 for "RRBP1 rewires cisplatin resistance in Oral Squamous Cell Carcinoma by regulating YAP-1"

### Supplementary Fig 2

A

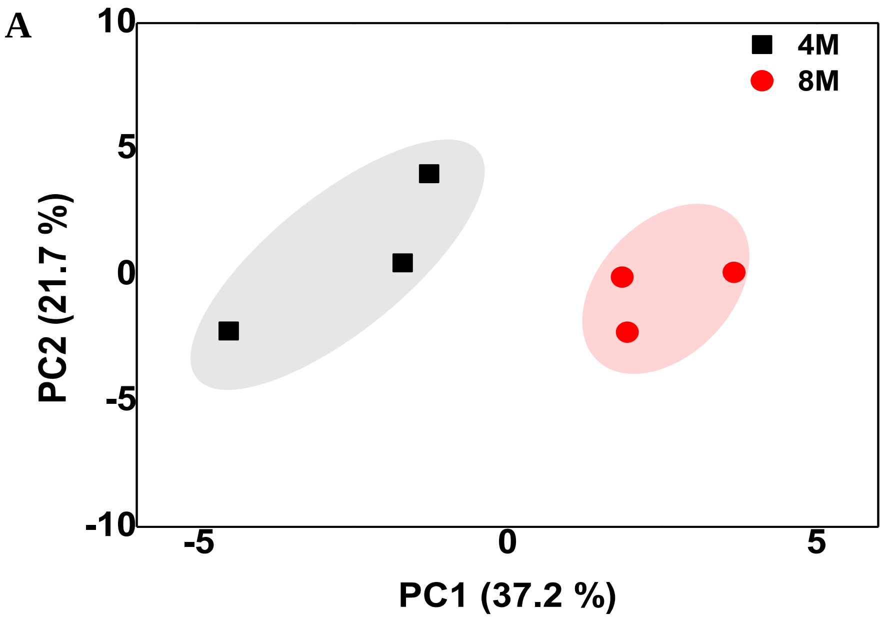


B

**RRBP1 (Ribosomal binding protein-1) [D].ALNQATSQVES.[K]**

>tr|A1A5C5|A1A5C5_HUMAN RRBP1 protein OS=Homo sapiens OX=9606 GN=RRBP1 PE=2 SV=1
MVFNEGEAQRLIEILSEKAGIIQDTWHKATQKGDPVAILKRQLEEKEKLLATEQEDAAVAKSKLREL
NKEMAAEKAKAAAGEAKVKKQLVAREQEITAVQARMQASYREHVKEVQQLQGKIRTLQEQLENG
PNTQLARLQQENSILRDALNQATSQVESKQNAELAKLRQELSKVSKELVEKSEAVRQDEQQRKALE
AKAAAFEKQVLQLQASHRESEEALQKRLDEVSRELCHTQSSHASLRADAEKAQEQQQMAELHS
KLQSSEAEVRSKCEELSGLHGQLQEARAENSQLTERIRSIEALLEAGQARDAQDVQASQAEADQQQ
TRLKELESQVSGLEKEAIELREAVEQQKVKNNDLREKNWKAMEALATAEQACKEKLHSLTQAKEE
SEKQLCLIEAQTMEALLALLPELSVLAQQNYTEWLQDLKEKGPTLLKHPPAPAEPSSDLASKLREAE
ETQSTLQAECDQYRSILAETEGMLRDLQKSVEEEEQVWRAKVGAAEEELQKSRVTVKHLEEIVEKL
KGELESSDQVREHTSHLEAELEKHMAAASAECQNYAKEVAGLRQLLLESQSQLDAAKSEAQKQSD
ELALVRQQLSEMKSHVEDGDIAGAPASSPEAPPAEQDPVQLKTQLEWTEAILEDEQTQRQKLTAEFE
EAQTSACLLQEELEKLRTAGPLESSETEEASQLKERLEKEKKLTSDLGRAATRLQELLKTTQEQLAR
EKDTVKKLQEQLEKAEDGSSSKEGTSV

C

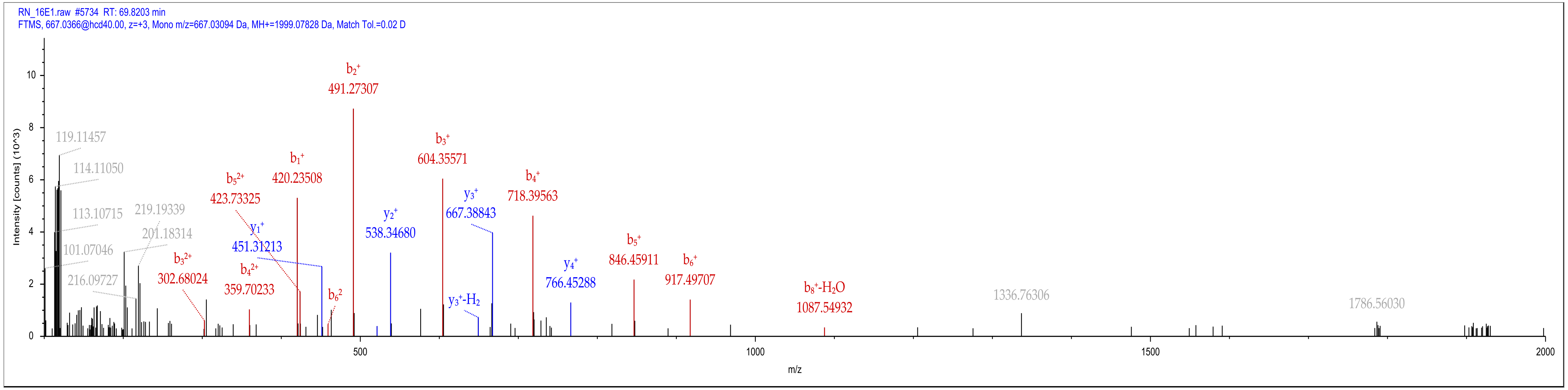


**Supplementary figure 2**: : **Fragmentation pattern of the peptide ALNQATSQVES (residues 149 to159) from the protein RRBP1 human**. A) peptide sequence of RRBP1 B) fragmentation of peptide **ALNQATSQVES**
