## Supplementary Figure 3 for "RRBP1 rewires cisplatin resistance in Oral Squamous Cell Carcinoma by regulating YAP-1"

### Supplementary data 3

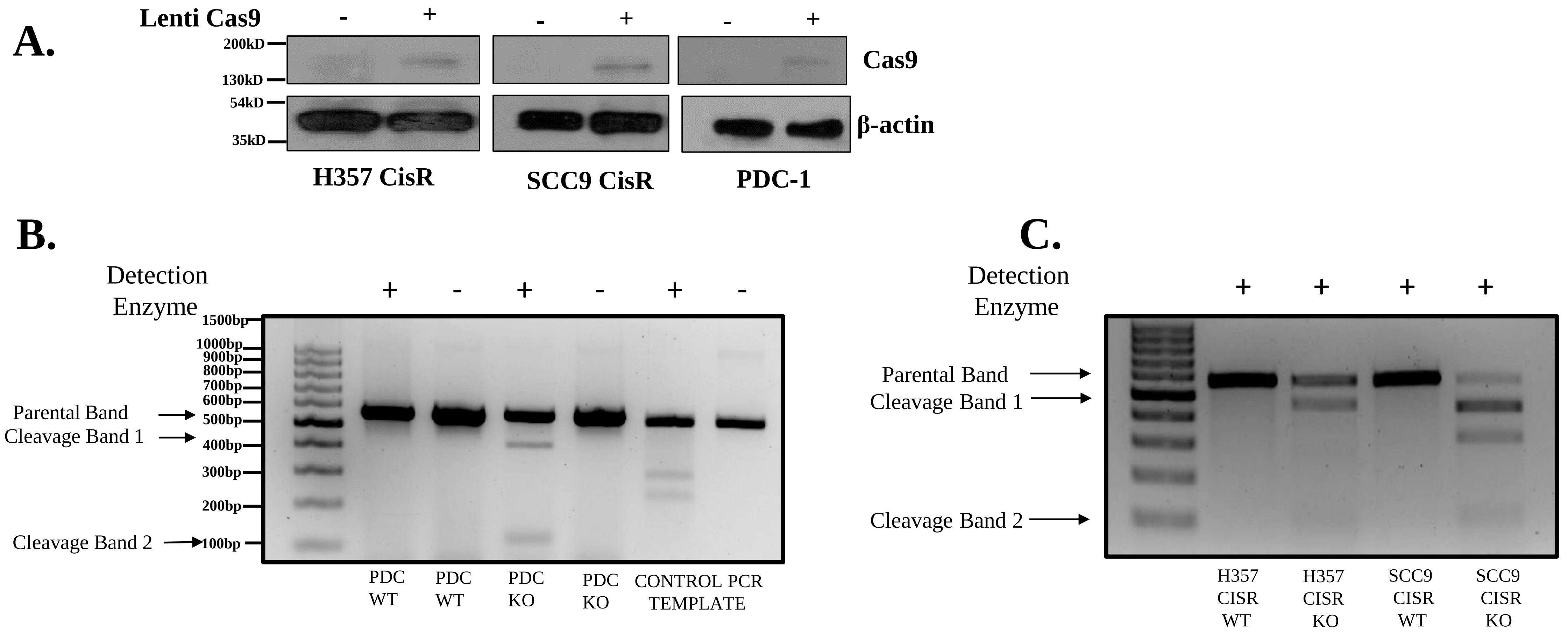


**Supplementary figure 3: The confirmation of RRBP1 KO in OSCC Chemoresistance cell line done by using GeneArt®Genomic Cleavage Detection (GCD) Kit (Life technology) with PCR primers listed in supplementary table** . A) Stable clones of CAS9 expressing chemoresistant OSCC lines were established as described in method section and lysates were isolated to perform immunoblotting against Cas9 and β-actin B) Agarose gel showing GCD assay to detect the on-target of RRBP1 targeting sgRNA in the indicated cell line in Cas9 overexpressed. For +ve control we used HPRT template that was provided in the kit.
