## Supplementary Figure 4 for "RRBP1 rewires cisplatin resistance in Oral Squamous Cell Carcinoma by regulating YAP-1"

### Supplementary Fig 4

A

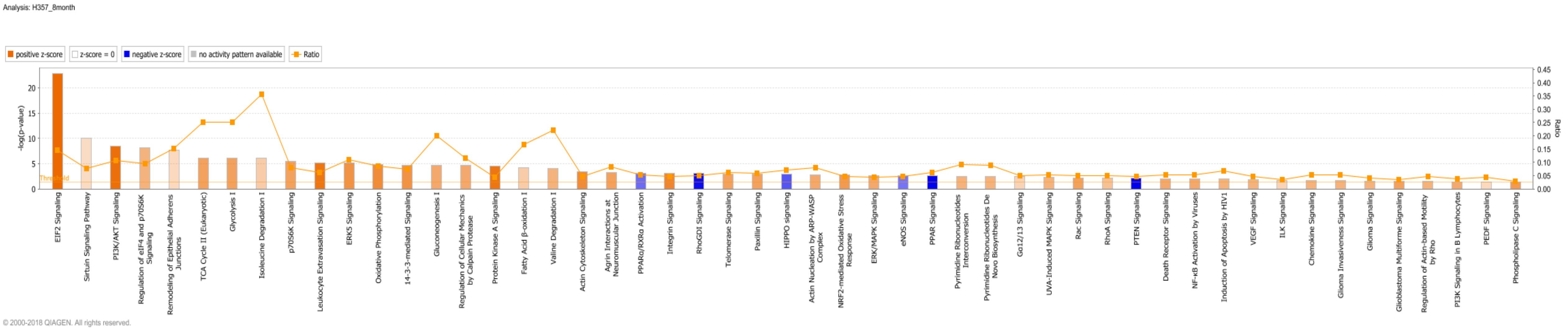


B

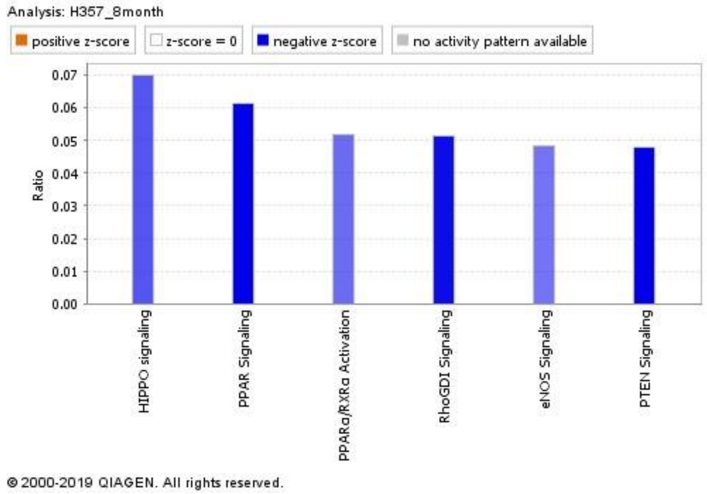


**Supplementary figure 4: Analysis of pathways involved in early and late cisplatin resistant OSCC lines:** The deregulated proteins, identified from global proteomics analysis, were converted to gene list and a functional analysis was carried out using Ingenuity Pathway Analysis**. A)** Top upregulated and downregulated canonical pathways in early vs. late cisplatin resistance line normalized with sensitive and list of pathways involved are indicated as bar diagram **B)** Top 6 down regulated pathway were selected among all pathway
