## Supplementary table 1 for "RRBP1 rewires cisplatin resistance in Oral Squamous Cell Carcinoma by regulating YAP-1"

| **Sl No** | **Tumor samples** | **Age/Sex** | **Site of disease** | **Clinical stage** |
| --- | --- | --- | --- | --- |
| **1** | **Patient#1** | 32/M | Tongue Lt lateral border | T4N1M0 |
| **2** | **Patient#2** | 29/F | Rt- Upper gingivobuccal maxillary antrum | T2N1Mx |
| **3** | **Patient#3** | 60/F | Upper alveolus | T4aN1M0 |
| **4** | **Patient#4** | 42/M | Tongue Lt lateral border | T4N1M0 |
| **5** | **Patient#5** | 60/M | Lt- Buccal mucosa | T4aN1M |
| **6** | **Patient#6** | 67/M | Tongue Lt lateral border | T4aN1Mx |
| **7** | **Patient#7** | 33/M | Tongue Lt lateral border | T3N1Mx |
| **8** | **Patient#8** | 55/F | Rt- Buccal mucosa | T4bN2bM0 |
| **9** | **Patient#9** | 50/M | Rt- Buccal mucosa | T4aN2bM0 |
| **10** | **Patient#10** | 42/M | Tongue Rt lateral border | T4aN2bM0 |
| **11** | **Patient#11** | 52/M | Tongue Lt lateral border | T2N1Mx |
| **12** | **Patient#12** | 38/M | Lt-Buccal Mucosa | T4N1M0 |
| **13** | **Patient#13** | 75/M | Rt- Buccal mucosa | T4aN2bM0 |
| **14** | **Patient#14** | 74/M | Tongue lateral border | T2N0M0 |
| **15** | **Patient#15** | 53/M | Tongue Lt lateral border | T4N0M0 |
| **16** | **Patient#16** | 34/M | Lt Lt-Buccal Mucosa | T4aN2bMx |
| **17** | **Patient#17** | 40/M | Tounge- lateral border | T2N2cM0 |
| **18** | **Patient#18** | 74/M | Tounge | T2N0M0 |
| **19** | **Patient#19** | 53/M | Oral cavity | T4N0M0 |
| **20** | **Patient#20** | 45/M | Tounge | T4aN2cM0 |
| **21** | **Patient#21** | 46/M | Tounge | T3N1M0 |
| **22** | **Patient#22** | 35/M | Tounge | T4aN2eM0 |
| **23** | **Patient#23** | 36/M | Tounge | cT4aN2aM0  cT4aN2cM0 |
| **24** | **Patient#24** | 48/M | Left Buccal Mucosa | T3N1Mx |
| **25** | **Patient#25** | 38/M | SCC Base of Tongue with extension into Oropharynx | T4aN2cMO |
| **26** | **Patient#26** | 36/M | Right Buccal Mucosa | T4aN2aM0 |
| **27** | **Patient#27** | 34/M | Left Buccal Mucosa | T4aN2bMx |
| **28** | **Patient#28** | 38/M | Right Gingiva buccal sulcus | T4N1Mx |
| **29** | **Patient#29** | 40/M | Tounge | T2N2M0 |

**Table-1: chemotherapy-naïve Patient details**

**Lt-Left,**

**Rt-Right**
