## Supplementary table 2 for "RRBP1 rewires cisplatin resistance in Oral Squamous Cell Carcinoma by regulating YAP-1"

| **Sl No** | **Tumor samples** | **Age**  **/Sex** | **Site of disease** | **Clinical stage** | **Chemotherapy (NACT)** | **Cycle** | **Response** |
| --- | --- | --- | --- | --- | --- | --- | --- |
| **1** | **Patient# 1**  **(PDC#1)** | 76/M | Tongue Rt lateral border | T4N0M0 | Paclitaxel + Cisplatin | 3 | non responder |
| **2** | **Patient#2** | 51/M | Rt- Buccal mucosa | T2N2bM0 | Docetaxel + Cisplatin+ 5FU | 2 | non responder |
| **3** | **Patient# 3** | 60/M | Tongue Rt lateral border | T3N1M0 | Paclitaxel + Cisplatin +5FU | 3 | non responder |
| **4** | **Patient#4** | 33/M | Rt- Lower Alveolar mucosa | T3N1Mx | Docetaxel + Cisplatin | 3 | non responder |
| **5** | **Patient#5** | 60/F | Tongue Lt lateral border | T4N0M0 | Docetaxel + Cisplatin+ 5FU | 3 | non responder |
| **6** | **Patient#6** | 59/M | Tongue Rt lateral border | T4aN1M0 | Docetaxel + Cisplatin+ 5FU | 3 | non responder |
| **7** | **Patient#7** | 46/M | Tongue | T4N3M0 | Docetaxel + Cisplatin+ 5FU | 3 | non responder |
| **8** | **Patient#8** | 55/F | Rt- Buccal Mucosa | T4aN2M0 | Docetaxel + Cisplatin+ 5FU | 2 | *Partial responder |
| **9** | **Patient#9** | 37/M | Tongue | T4N3M0 | Docetaxel + Cisplatin+ 5FU | 2 | non responder |
| **10** | **Patient#10** | 27/M | Lt-Buccal Mucosa | T4N2M0 | Docetaxel + Cisplatin+ 5FU | 2 | *Partial responder |
| **11** | **Patient#11** | 46/F | Rt- oral cavity | T4N1M0 | Docetaxel + Cisplatin+ 5FU | 3 | non responder |
| **12** | **Patient#12** | 42/M | Rt- Buccal Mucosa | TxN3bM0 | Paclitaxel + Cisplatin+ 5FU | 2 | non responder |
| **13** | **Patient#13** | 30/M | Tongue Rt lateral border | T2N0Mx | Paclitaxel + Cisplatin+ 5FU | 3 | non responder |
| **14** | **Patient#14** | 52/M | Rt- Buccal Mucosa | T4N2M0 | Docetaxel + Cisplatin+ 5FU | 3 | non responder |
| **15** | **Patient#15** | 32/M | Tongue | T3N1M0 | Docetaxel + Cisplatin+ 5FU | 3 | non responder |
| **16** | **Patient#16** | 35/M | Tongue | T4aN2aM0 | Docetaxel + Cisplatin+ 5FU | 3 | *Partial responder |
| **17** | **Patient#17** | 36/M | Left Buccal Mucosa | T4aN2aM0 | Docetaxel + Cisplatin+ 5FU | 3 | non responder |
| **18** | **Patient #18** | 38/M |  | T4bN2bMO | Docetaxel + Cisplatin+ 5FU | 2 | *Partial responder |
| **19** | **Patient # 19** | 55/M | Tounge, Left lateral border, |  | Docetaxel + Cisplatin+ 5FU | 3 | *Partial responder |
| **20** | **Patient # 20** | 35/M | Tounge | T4aN2eM0+ | Docetaxel + Cisplatin+ 5FU | 3 | non-Responder |
| **21** | **Patient # 21** | 36/M | Left Buccal Mucosa | cT4aN2aM0 | Docetaxel + Cisplatin+ 5FU | 3 | Non Responde |
| **22** | **Patient # 22** | 39/M | Right mandible | cT4bN0Mx | Docetaxel + Cisplatin+ 5FU |  | Non Responder |
| **23** | **Patient # 23** | 55/M | Tounge, Left lateral border, | cT4aN2cM0 | Docetaxel + Cisplatin+ 5FU | 3 | Non responder |

**Table-2: Chemotherapy-non-responders patient Detail**

***Partial response-** Patient initially responded to chemotherapy but after 1-2 cycles became non responded.

**Chemotherapy Doses**: **Cisplatin**: 100mg. **Paclitaxe**l: 260 mg, **Docetaxel**: 100mg, **5FU**:1000mg
