## Supplementary table 3 for "RRBP1 rewires cisplatin resistance in Oral Squamous Cell Carcinoma by regulating YAP-1"

**Table 3: List of primer used in this study**

| Sr No. | Primer Set | primer | Application |
|---|---|---|---|
| 1 | **RRBP1 F** | GGAGGAAGTACAGCCAGCAAAG | qRT PCR |
| | **RRBP1 R** | CTGCCAATGCCATCGTAATCACTC | |
| 2 | **GAPDH F** | TCGGAGTCAACGGATTTGGT | qRT PCR |
| | **GAPDH R** | TTGCCATGGGTGGAATCATA | |
| 3 | **CCTGF F** | CAAGGGCCTCTTCTGTGACT | qRT PCR |
| | **CTGF R** | ACGTGCACTGGTACTTGCAG | |
| 4 | **CYR61 F** | CCTCGGCTGGTCAAAGTTAC | qRT PCR |
| | **CR61 R** | TTTCTCGTCAACTCCACCTC | |
| 5 | **JAG1 F** | GAAGCAGAACACGGGCGTT | qRT PCR |
| | **JAG1 R** | CAGGTCACGCGGATCTGAT | |
| 6 | **AXL 1 F** | CCAGGACACCCCAGAGGTGCTAAT | qRT PCR |
| | **AXL1 R** | TGGTGGACTGGCTGTGCTTGC | |
| 7 | **YAP-1 F** | CTCGAACCCCAGATGACTTC | qRT PCR |
| | **YAP-1 R** | CCAGGAATGGCTTCAAGGTA | |
| 8 | **RRBP1 GCD F** | GGAGATTCAGATGACCAGGG | RT PCR for genomic cleavage detection Assay |
| | **RRBP1 GCD R** | TCAGGCTGTACCTTGTTGAG | |
